# Metabolic responses to physiological stressors assessed using continuous glucose monitoring integrated with indirect calorimetry in mice

**DOI:** 10.64898/2026.07.13.738308

**Authors:** Evelyn Y. Zhou, Corey D. Holman, Michelle Lee, William B. Rubio, Ryan Calhoun, Qingwei Chu, Joseph A. Baur, Alexander S. Banks

## Abstract

Continuous glucose monitoring (CGM) in rodents has provided unprecedented temporal resolution of glycemic dynamics in vivo. Even in the absence of deliberate perturbation, glucose levels in mice are dynamic, fluctuating in response to the timing and duration of feeding events, changes in neurological and hormonal states, physical activity, and photoperiod. To obtain a comprehensive view of metabolic adaptations under common experimental conditions, we monitored freely moving mice simultaneously using CGM and indirect calorimetry to quantify glucose, food intake, physical activity and metabolic rate. We characterized glycemic and metabolic responses to routine laboratory interventions, including short-term and overnight fasting, refeeding, tail blood sampling during glucose tolerance tests, changes in ambient temperature to cold or thermoneutral conditions, and access to running wheels.

We found that food removal induced a robust, transient stress response characterized by increased blood glucose, body temperature, energy expenditure, and physical activity. However, prolonged fasting ultimately led to hypoglycemia and torpor. The magnitude and variability of glycemic responses to insulin tolerance tests were strongly influenced by fasting duration, and tail-tip blood collection itself elicited substantial hyperglycemia. In contrast to prolonged fasting, refeeding produced relatively modest and transient effects on glucose and energy expenditure. Cold exposure elicited increased energy expenditure along with a sustained hyperglycemic response. Voluntary wheel running induced transient increases in glucose and metabolic activity and promoted a shift toward increased fatty acid oxidation. Together, these findings demonstrate that common laboratory manipulations exert dynamic, often substantial effects on glycemia and whole-body metabolism that are readily revealed by CGM and indirect calorimetry.

## Introduction

Assessments of metabolic phenotypes routinely include indirect calorimetry to measure physiological parameters such as energy expenditure, alongside standard assays such as glucose tolerance tests (GTTs) and insulin tolerance tests (ITTs) to evaluate systemic glycemic control. However, these methods are usually conducted separately, as GTTs and ITTs require intermittent blood sampling with hand-held glucometers and associated animal handling, which interrupts indirect calorimetry recordings. Continuous glucose monitoring (CGM) has emerged as a powerful tool for rodent studies, enabling minimally invasive, high-resolution tracking of glucose dynamics over time, reducing handling and capturing rapid fluctuations that are missed by conventional sampling (1). When combined with indirect calorimetry, CGM allows for simultaneous assessment of glucose, energy expenditure, respiratory exchange ratio (RER), and activity, providing a comprehensive view of whole-body metabolic physiology (2).

Despite these advances, stress remains a critical and often underappreciated confounder in metabolic studies. Stress modulates glucose metabolism primarily through activation of the sympathetic nervous system and hypothalamic-pituitary-adrenal (HPA) axis, leading to rapid increases in circulating glucose (3,4). In mice, common experimental manipulations such as fasting, handling, and changing environmental temperatures can elicit acute physiological responses that are often difficult to disentangle from the metabolic processes under investigation. For example, while fasting is routinely used to standardize baseline glucose levels (5), it also alters energy balance, modulates hormone secretion, and increases intragroup variability of GTT results (6,7). Similarly, repeated blood sampling via tail-tip incision and restraint has been shown to elevate stress hormones and influence glucose dynamics (8). Environmental temperature further modulates metabolic physiology in mice, which are typically housed below thermoneutrality. Sub-thermoneutral housing conditions require substantial energy expenditure and substrate utilization to maintain core body temperature (9,10).

In this study, we combined CGM and indirect calorimetry to concurrently characterize the impact of commonly used experimental conditions on glucose homeostasis and whole-body physiology in mice. By systematically examining the effects of fasting duration, handling-associated blood sampling, ambient temperature changes, and voluntary exercise, we provide a high-resolution framework for understanding how experimental context influences metabolic outcomes and highlight the importance of minimizing disturbances in metabolic studies.

## Results

### Fasting-induced stress response and acute metabolic adaptations

In rodent metabolic studies, fasting is commonly performed to standardize baseline conditions prior to metabolic assays such as GTT and ITT (11,12). However, fasting itself is a physical stressor that can independently alter metabolic parameters (6). We therefore examined the physiological response of mice to short-term fasting of 2, 4, or 6-hour duration, compared to mice with ad libitum access to food. Mice were simultaneously monitored by CGM and indirect calorimetry to assess blood glucose, body temperature, energy expenditure (EE), respiratory exchange ratio (RER), physical activity, and food intake.

Removal of food triggered a rapid, transient glycemic increase, consistent with an acute stress response and characterized by coordinated elevations in blood glucose, body temperature, EE, and physical activity relative to ad libitum-fed controls (Figure 1A). This apparent stress response resolved within approximately 2 hours of food removal, after which these parameters returned to levels comparable to controls. By the final hour of the fasting period, there were no significant differences among groups in any parameter (Figure 1B), indicating that short-term fasting induces only transient metabolic perturbations in a subset of measured physiological parameters.

**Figure 1.**
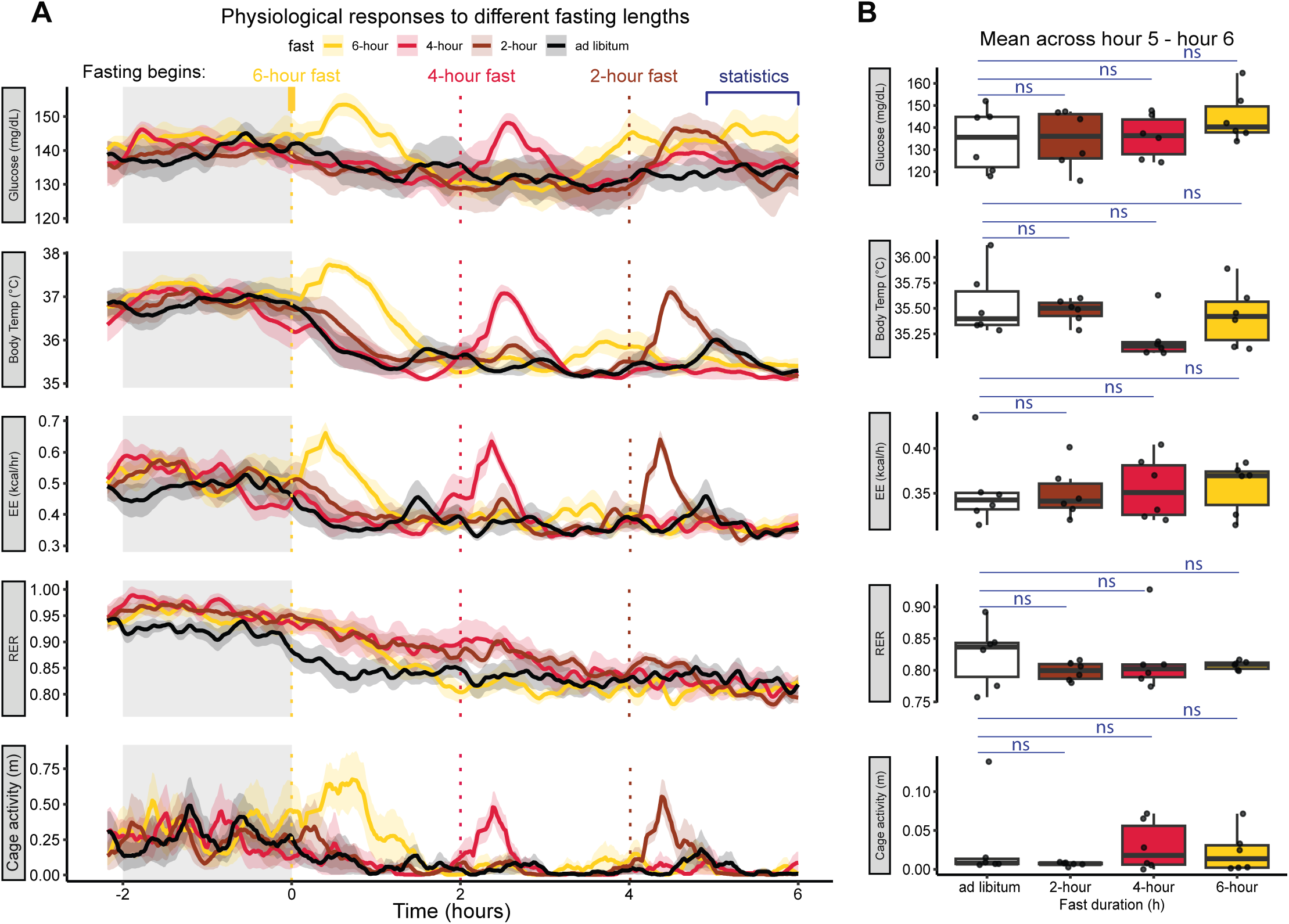
Physiological responses during fasting (n=6). **A**) Continuous glucose monitoring and indirect calorimetry recordings in mice over a 6-hour window, with fasting initiated at staggered time points to achieve different fasting durations. Grey shadings indicate the dark photoperiod. **B**) Boxplots showing mean values over hour 5–6 for each parameter. Points represent individual mice. Statistics by posthoc; ns, not significant.

Notably, EE and physical activity began to increase before food removal. Since fasting was initiated by removing the food containers in the indirect calorimetry system, the presence of the investigator and cage disturbance may have caused an anticipatory stress response in the mice.

### Overnight fasting, torpor, and refeeding responses

We next assessed the impact of a more severe overnight (16 hr) fast and subsequent refeeding on glucose homeostasis and whole-body metabolism. Food access was removed 2 hours before the onset of the dark photoperiod by closing food access control doors in the indirect calorimetry system. As with shorter fasts, the initial phase of overnight fasting was accompanied by a small transient increase in blood glucose, body temperature, EE, and physical activity upon food access removal, and a second increase in these parameters coincided with the start of the dark photoperiod (Figure 2A).

**Figure 2.**
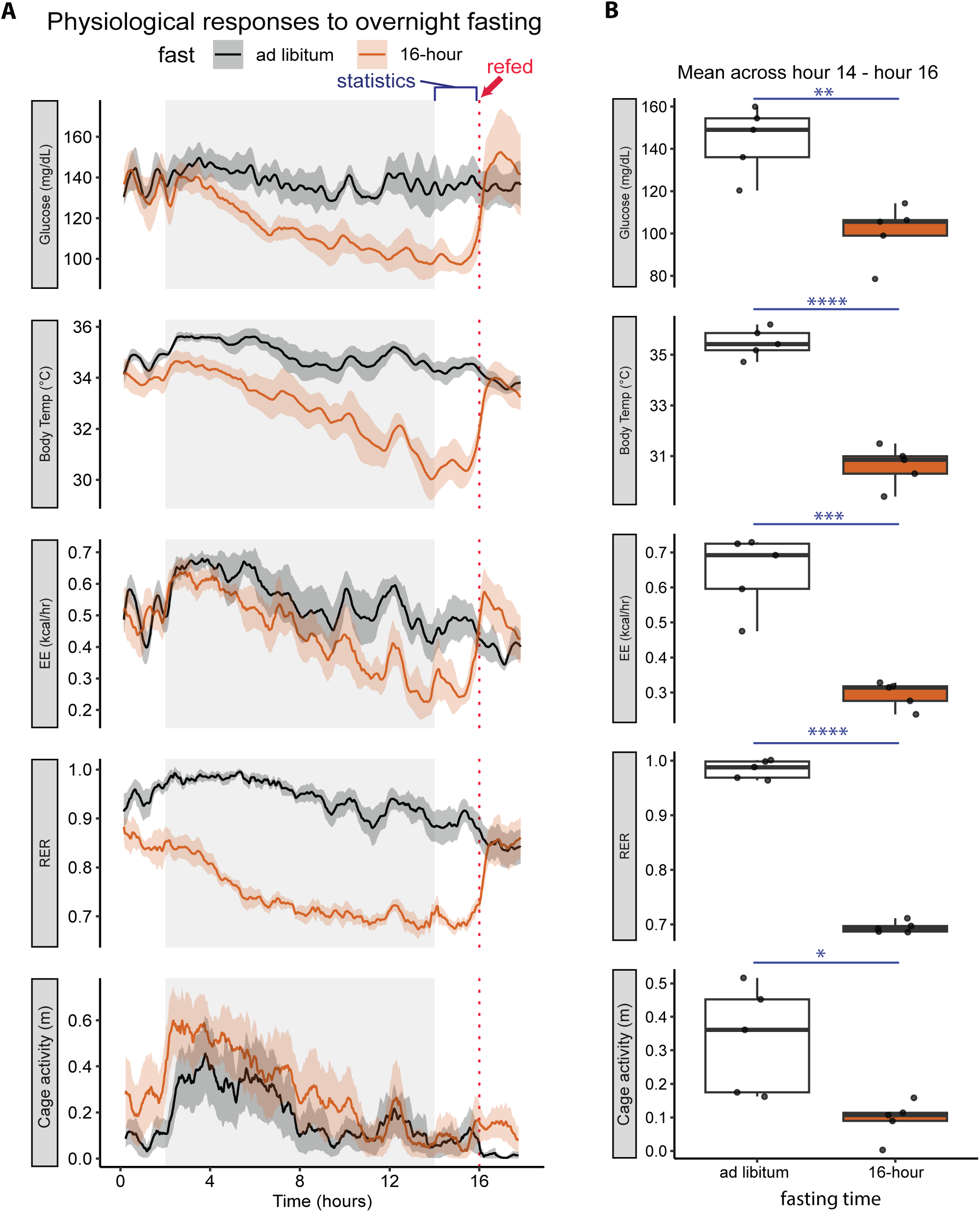
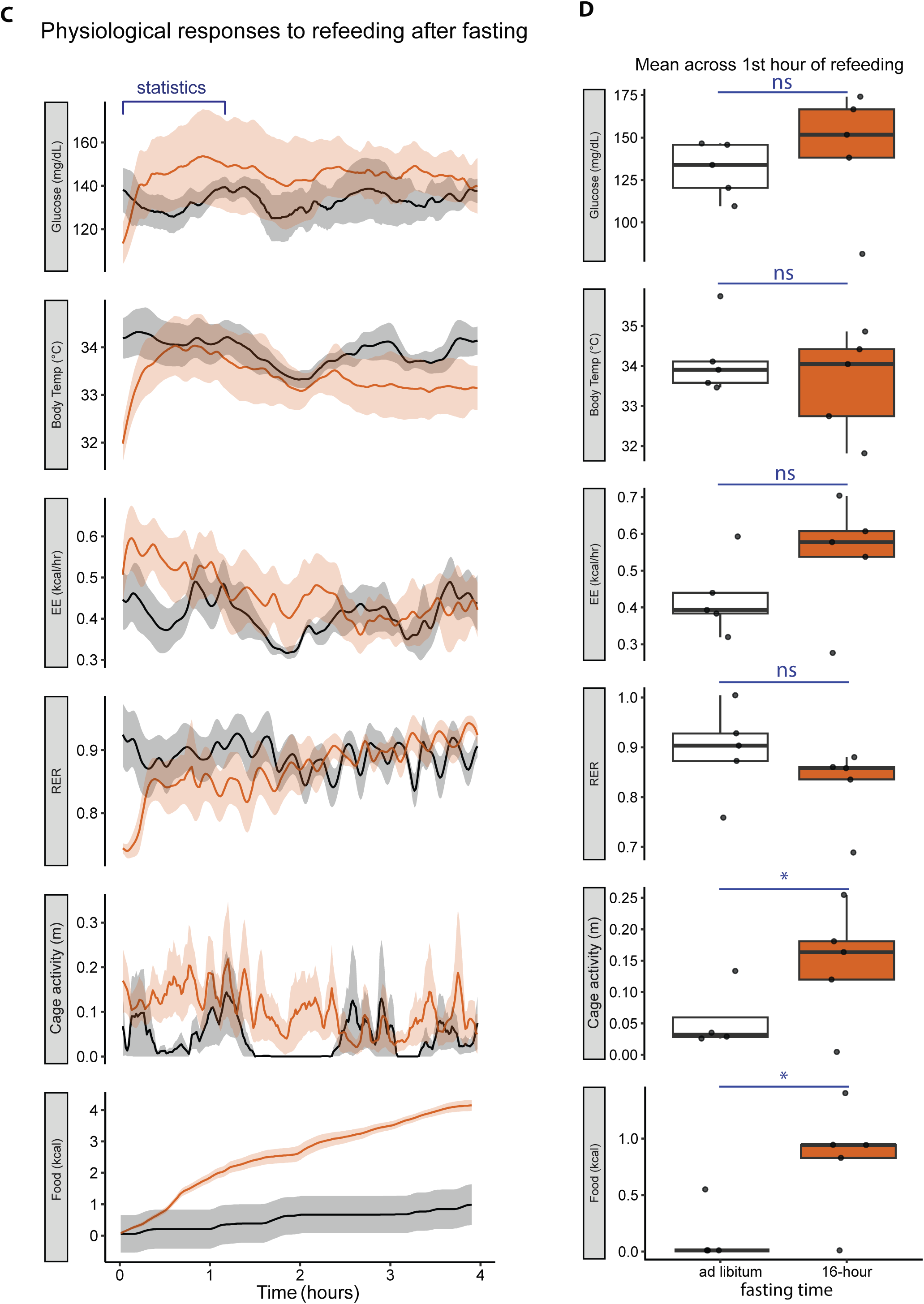
Effects of overnight fasting and refeeding on metabolic parameters in chow-fed mice (n=5). **A)** Recordings of chow-fed mice during 16h overnight fasting and 4h refeed. Solid lines represent the mean values at each minute; shaded areas represent SEM, and dotted lines represent the start of refeed. Grey shadings indicate the dark photoperiod. **B)** Boxplots showing mean values calculated over hours 14–16. Each point represents one mouse. Statistics by two-tailed unpaired t tests. **C)** Recordings of refeeding following overnight fast, mice were refed at t = 0. Solid lines represent the mean value at each minute, shaded areas represent SEM. **D)** Boxplots showing mean values calculated over hours 0-1 after refeeding. Each point represents one mouse. Statistics by two-tailed unpaired t tests. *p<0.05, **p<0.01, ***p<0.001, ****p<0.0001.

However, once fasting extended into the dark photoperiod, mice developed progressive hypoglycemia, along with significant reductions in body temperature and RER, indicating a marked shift toward lipid oxidation. By the final two hours of the fast, all measured metabolic parameters, including glucose, body temperature, EE, RER, and physical activity were significantly altered compared to ad libitum controls (Figure 2B). These changes coincided with behavioral and physiological features consistent with torpor, which was observed in mice fed either standard chow (Figure 2A, 2B) or a high-fat diet (HFD, Figure S1).

Refeeding caused a rapid increase in blood glucose, body temperature, energy expenditure, RER, and physical activity, normalizing most parameters within the first hour (Figure 2C, 2D). Unsurprisingly, food intake was significantly greater in refed mice compared to unfasted controls. These results indicate that refeeding elicited a rapid metabolic response responsible for normalizing the profound alterations induced by overnight fasting and torpor.

### Effect of fasting duration on insulin tolerance

Although glucose levels converged to ad libitum levels by the final hour of 2-, 4-, and 6-hour fasts (Figure 1), it remained unclear whether fasting duration would influence the glycemic response to an insulin bolus. To address this, we performed insulin tolerance tests (ITTs) in mice fasted for 2, 4, or 6 hours, as well as in mice with ad libitum access to food. Despite similar baseline metabolic parameters immediately before insulin treatment, glycemic excursions during the ITT varied markedly with fasting duration. Based on Area of curve (AOC) for blood glucose, mice tested under ad libitum-fed conditions exhibited the smallest insulin-mediated decrease, whereas 6-hour-fasted mice displayed the greatest reduction in glycemia. Intermediate responses were observed in the 2- and 4-hour fasted groups (Figure 3A, 3B). The absolute between-animal variability in the integrated glycemic response, measured as the group standard deviation (SD) of AOC, increased with fasting duration, consistent with the progressively larger insulin-induced decreases in blood glucose. Ad libitum-fed mice showed the smallest SD in insulin-induced glucose lowering, whereas 6-hour-fasted mice showed the largest variability. These data indicate that both the magnitude and variability of ITT outcomes are highly sensitive to pre-test fasting duration, even when other metabolic parameters appear similar at the end of each respective fast (Figure 1). Similar results were observed for animals on an HFD (Figure S2).

**Figure 3.**
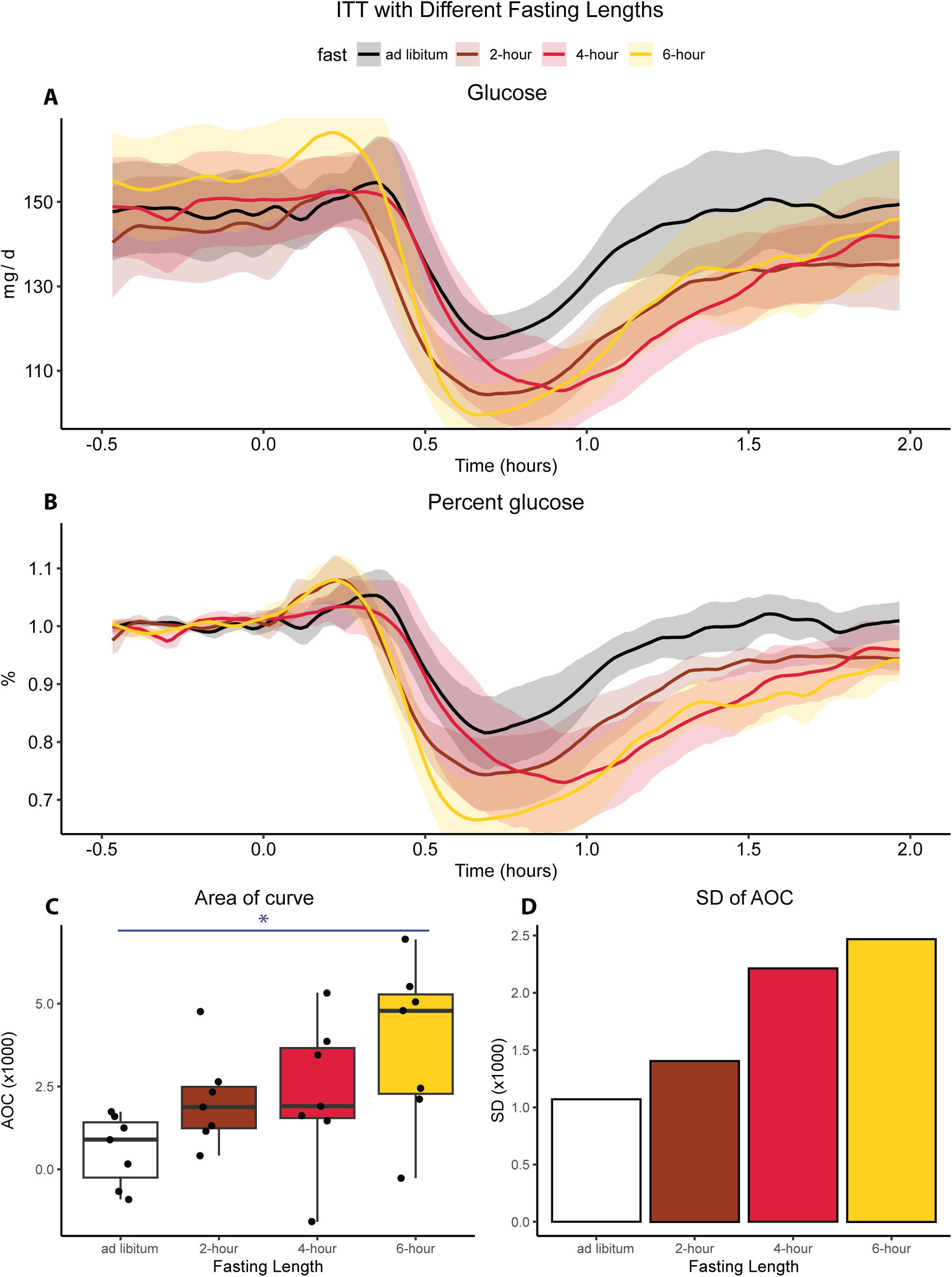
Effects of fasting duration on insulin tolerance test in chow-fed mice (n=7). **A)** Continuous glucose monitoring during ITTs plotted in absolute experimental time. Mice under-went no fasting (0-hour), 2-, 4-, or 6-h fasting prior to the insulin injection (t = 0). **B)** Glucose values expressed as percent of baseline for the same recordings. Baseline glucose was defined as the average glucose during the 1 h preceding insulin injection. **C)** Area of the curve (AOC) calculated from percent glucose traces. Each point represents one mouse. Statistical significance was detemined by one-way ANOVA; *p<0.05. **D)** Standard deviation (SD) of AOC across mice for each fasting duration.

### Effects of tail blood sampling on glucose tolerance test

In addition to fasting, routine blood glucose measurements using hand-held glucometers in metabolic assays introduce another physical stressor, as they require handling and tail-tip sampling (8). To determine how tail blood sampling influences metabolic readouts during glucose tolerance tests (GTTs), we performed GTTs in mice implanted with CGM. In the first condition, control mice received a single intraperitoneal (i.p.) glucose injection and were monitored continuously for glucose and body temperature using noninvasive telemetry, without tail blood sampling. Under these conditions of a CGM-GTT, chow-fed mice exhibited a relatively modest glycemic excursion, whereas mice maintained on an HFD showed a larger increase in blood glucose (Figure 4A). We then performed a conventional GTT on the same CGM-implanted mice, in which blood glucose was also measured by handheld glucometer via tail-tip sampling at 0, 15, 30, 60, 90, and 120 minutes following i.p. glucose injection. Glucose levels were also recorded simultaneously by CGM, and glucometer-derived samples from tail blood matched well to CGM data, indicating adequate CGM calibration. In chow-fed mice, the act of repeated tail sampling induced an exaggerated glycemic excursion with a mean AOC 3.2-fold greater than in the CGM-GTT. In HFD-fed mice, tail sampling further increased the hyperglycemic response to an AOC 2.7-fold greater than in the CGM-GTT. Furthermore, the tail sampling in both diet conditions resulted in a more gradual fall in blood glucose following the glucose peak compared to the sharp concave fall in blood glucose of control mice. These findings indicate that the stress of tail blood collection is a major, often unrecognized, contributor to glycemic excursions during standard GTT protocols (Figure 4B).

**Figure 4.**
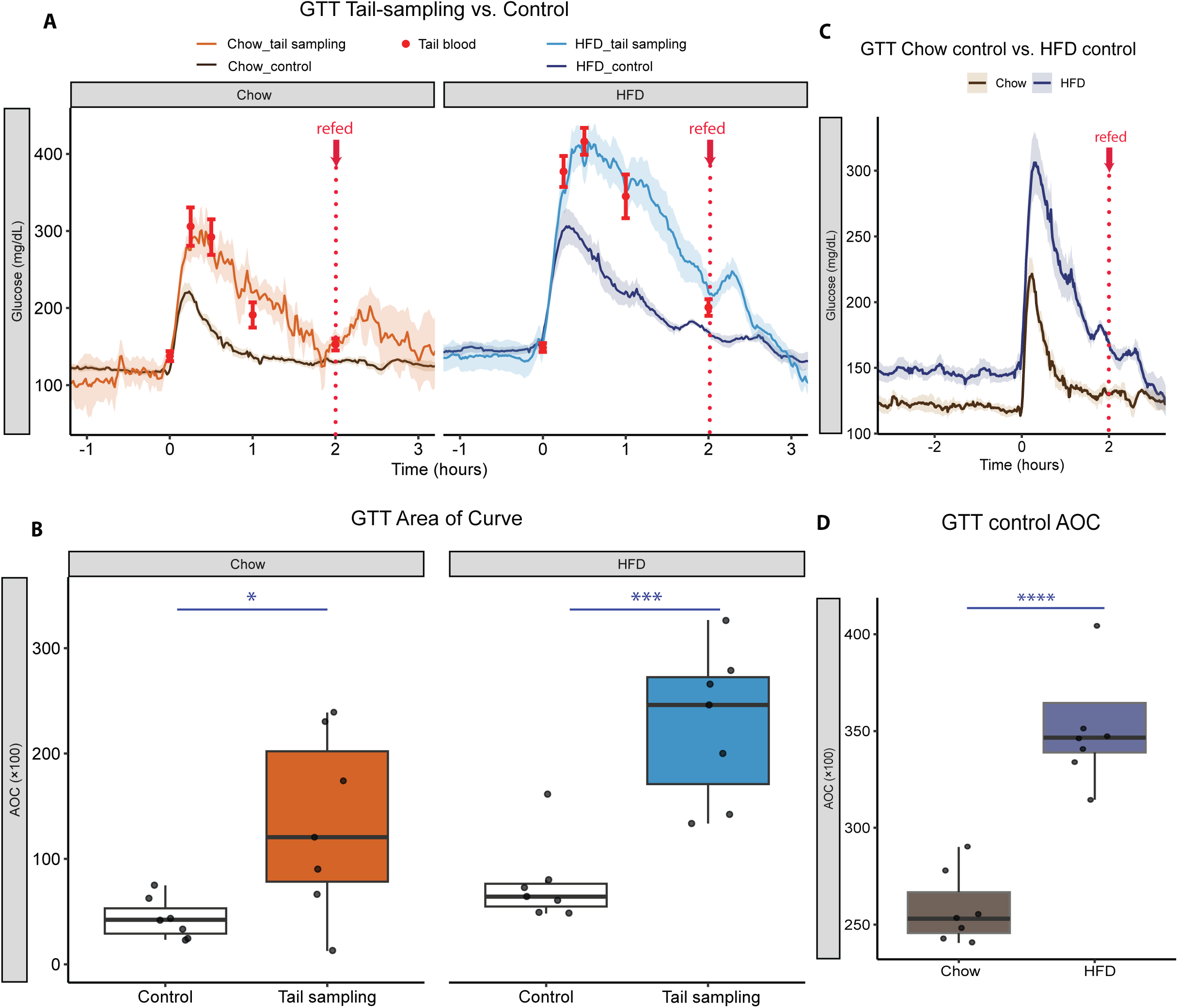
Effects of tail blood sampling on GTT and ITT in chow- and HFD-fed mice (n=7). **A)** glucose tolerance tests (GTT) with continuous glucose monitoring comparing without (control) and with tail blood sampling in chow- and high-fat diet (HFD)–fed mice. **B)** Area of the GTT glucose excursion curves. **C)** Comparison of chow and HFD control CGM GTT. **D)** AOC for GTT for chow vs HFD. Glucose injections at t = 0. Shaded areas indicate SEM. Red points represent blood glucose measurements obtained by glucometer from tail sampling Each point represents one mouse. Statistics by two-tailed unpaired t tests *p<0.05, **p<0.01, ***p<0.001.

We next directly compared GTT results in chow and HFD mice monitored solely with CGM. HFD mice exhibited higher baseline glucose levels. Using AOC, which accounts for different baseline glycemia, we observed a significantly greater AOC in HFD mice during the GTT compared to chow-fed mice (Figure 4C, 4D), consistent with impaired glucose clearance and reduced insulin sensitivity in HFD-fed mice.

### Glycemic response to changes in ambient temperature

In addition to physical stressors such as fasting and handling, environmental temperature is another key factor that can influence metabolic physiology in mice. Rodents respond to cold exposure by increasing thermogenesis and EE, but the accompanying effects on glycemia are less well characterized. We therefore investigated how acute changes in environmental temperature affect blood glucose dynamics. In a short-term intervention, we gradually decreased ambient temperature from 22°C to 6°C over a 2-hour period while monitoring mice with CGM and indirect calorimetry, then maintained 6°C for an acute cold exposure of 6 hours.

The acute 6-hr cold exposure elicited a robust thermogenic response in chow- and HFD-fed mice, characterized by increasing EE to maintain body temperature despite a significant reduction in core body temperature. In the chow-fed mice, we observed a transient increase in RER corresponding to an acute rise in food intake, then a gradual decrease in RER consistent with increased fatty acid oxidation used to fuel cold-induced thermogenesis. Concomitantly, we observed a nonsignificant trend toward elevated blood glucose (Figure 5A, 5B). In cold-challenged HFD mice, the rise in blood glucose was significantly greater than in HFD mice maintained at room temperature. The counterintuitive hyperglycemic response despite the increased energy demand observed in the HFD mice suggests that acute cold stress adaptively alters glucose homeostasis, likely through coordinated activation of sympathetic outflow and counterregulatory hormonal pathways. Besides an initial increase in physical activity during the two-hour cold ramp, there was no significant difference in physical activity in either group (Figure 5A, 5B).

**Figure 5.**
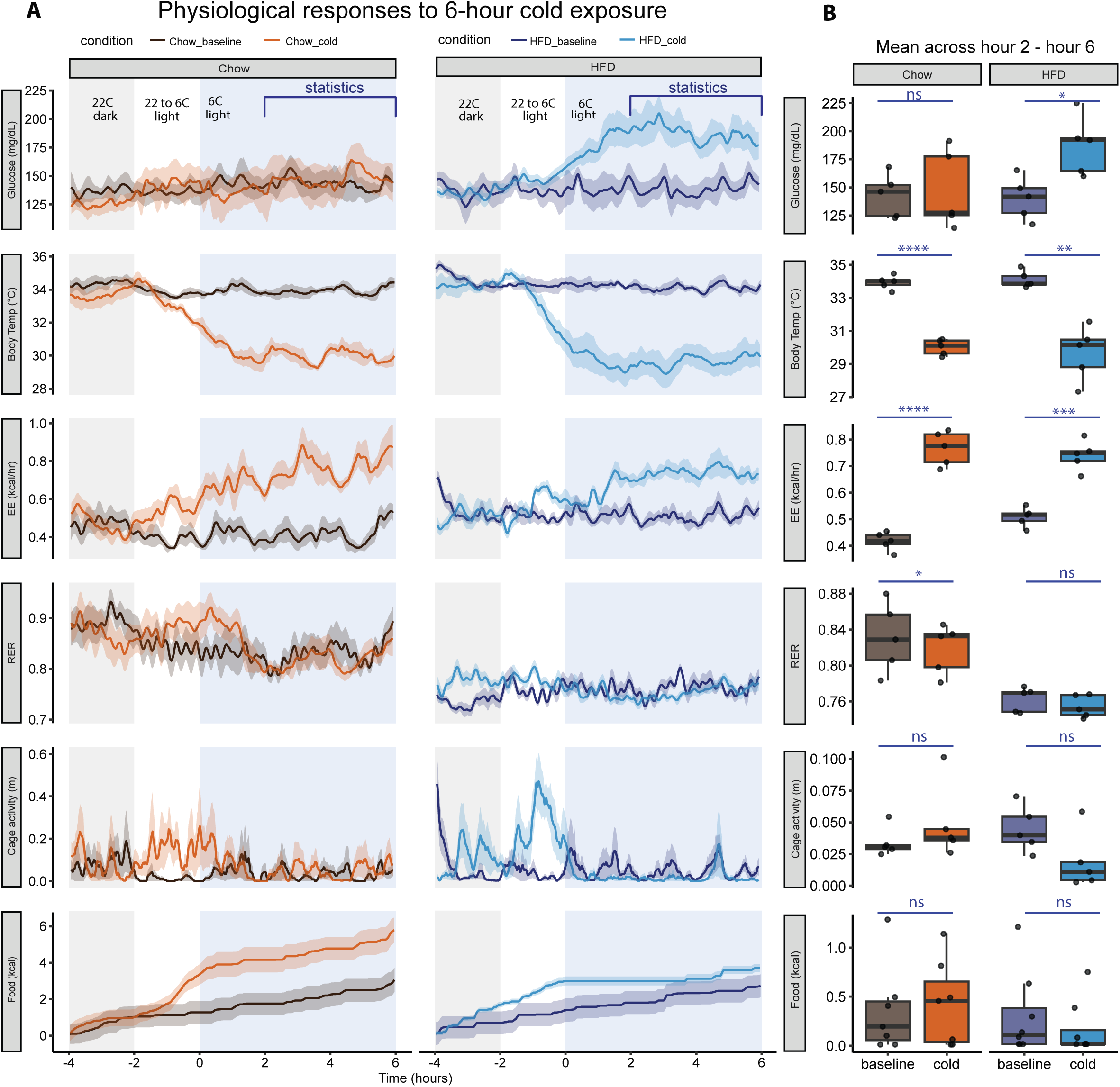
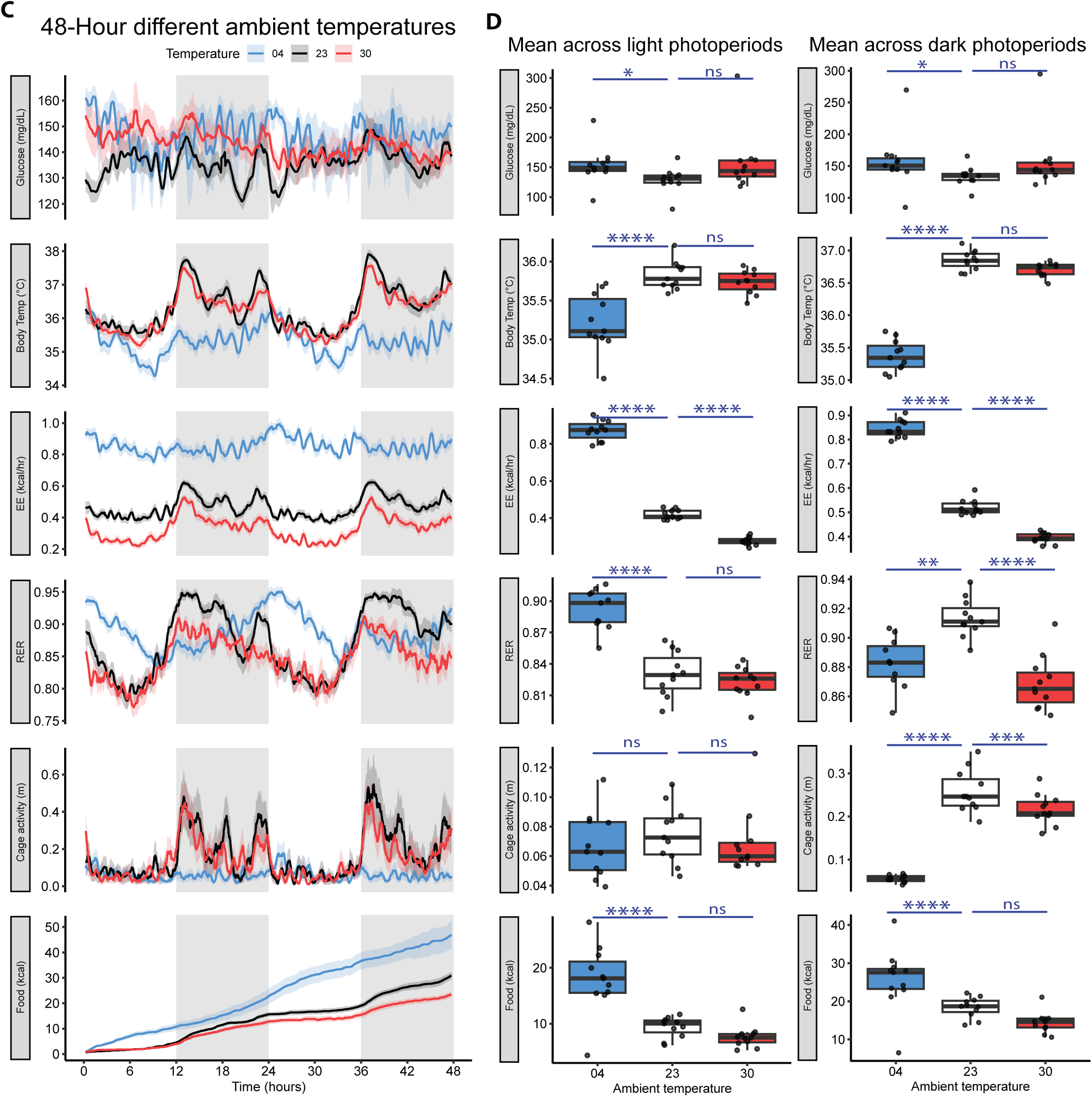
Metabolic responses to different temperatures in chow- and HFD-fed mice. **A)** Continuous recordings during a 6-h cold exposure (n=5). Mice were maintained at 22C from hour −4 to −2, ambient temperature began to decrease at hour −2 and reaches 4C at t = 0. Blue shadings indicate the cold period; grey shadings indicate the dark photoperiod. **B)** Boxplots showing mean values calculated over hours 2–6 of cold exposure. Each point represents one mouse. Statistics by two-tail unpaired t-test; ns, not significant, *p<0.05, **p<0.01, ***p<0.001, ****p<0.0001. **C)** Continuous recordings of chow-fed mice housed at different ambient temperatures (4°C, 23°C, and 30°C) over 48h (n=11). Gray shadings indicate dark photoperiods. Data are shown as mean ± SEM. **D)** Boxplots showing mean values calculated across light photoperiods and dark photoperiods. Statistics by post-hoc test. ns, not significant; *p<0.05, **p<0.01, ***p<0.001, ****p<0.0001.

To test whether cold-induced changes could be sustained past the 6-hr monitoring period, we performed CGM and indirect calorimetry monitoring for 48 hours on chow-fed mice housed at thermoneutrality (TN, 30°C), room temperature (RT, 23°C), or cold ambient temperatures (4°C). Glycemia was lowest in mice housed at room temperature but was significantly elevated during both light and dark photoperiods in mice housed at 4°C ambient temperature. Similarly, these cold-challenged animals defended significantly lower body temperatures and exhibited greater EE and food intake during both light and dark photoperiods. RER of 4°C-housed mice was lower than RT controls in the dark photoperiod and greater than RT controls in the light photoperiod. Physical activity was lower in cold-challenged mice in the dark photoperiod and not significantly different in the light photoperiod (Figure 5C, 5D, 5E). Mice housed at thermoneutrality had no significant difference in glucose or body temperature. However, they did exhibit significantly lower EE and dark phase RER compared to RT controls. In TN conditions, physical activity was decreased in the dark but not in the light. Conversely, food intake was decreased in the light but not in the dark (Figure 5C, 5D, 5E). These data demonstrate that changes in ambient temperature can alter glucose levels in C57BL/6J mice.

### Glycemic response to voluntary wheel running exercise

Finally, we examined the impact of voluntary exercise on glucose and other physiological parameters in mice given short-term, free access to a running wheel. While exercise is known to promote endogenous glucose production (13) in addition to driving glucose oxidation and heat generation, we observed no statistically significant change in glucose or body temperature on a group level. In contrast, RER exhibits a pronounced decrease in both light and dark photoperiods during wheel access, indicating a shift toward increased fatty acid oxidation (Figure 6A, 6B). Total EE was only modestly but not significantly increased, consistent with the prior findings suggesting that most of the energetic cost of exercise is offset by reducing the need for other means of thermogenesis (14). Unsurprisingly, voluntary wheel running led to a significant decrease in the dark photoperiod cage activity with these animals substituting wheel activity for cage ambulatory activity. Additionally, examination of individual mouse recordings revealed transient increases in glucose, body temperature, energy expenditure, and RER that coincided with bouts of wheel running (Figure 6C). These bout-associated increases likely reflect rapid, coordinated physiological responses to acute wheel-running activity, in which muscle contraction directly drives heat production while increased energetic demand promotes endogenous glucose production to support working muscle. The transient elevations in RER suggest a brief shift toward carbohydrate utilization, indicating that glucose is mobilized and utilized to meet the immediate energetic demands of activity. These dynamics are consistent with prior findings demonstrating that metabolic parameters fluctuate in ultradian cycles tightly linked to episodic activity bouts (15,16). Notably, these dynamic, within-animal responses were not apparent in the averaged data, suggesting that voluntary exercise in untrained animals induces short-lived metabolic changes that are masked when aggregated over time.

**Figure 6.**
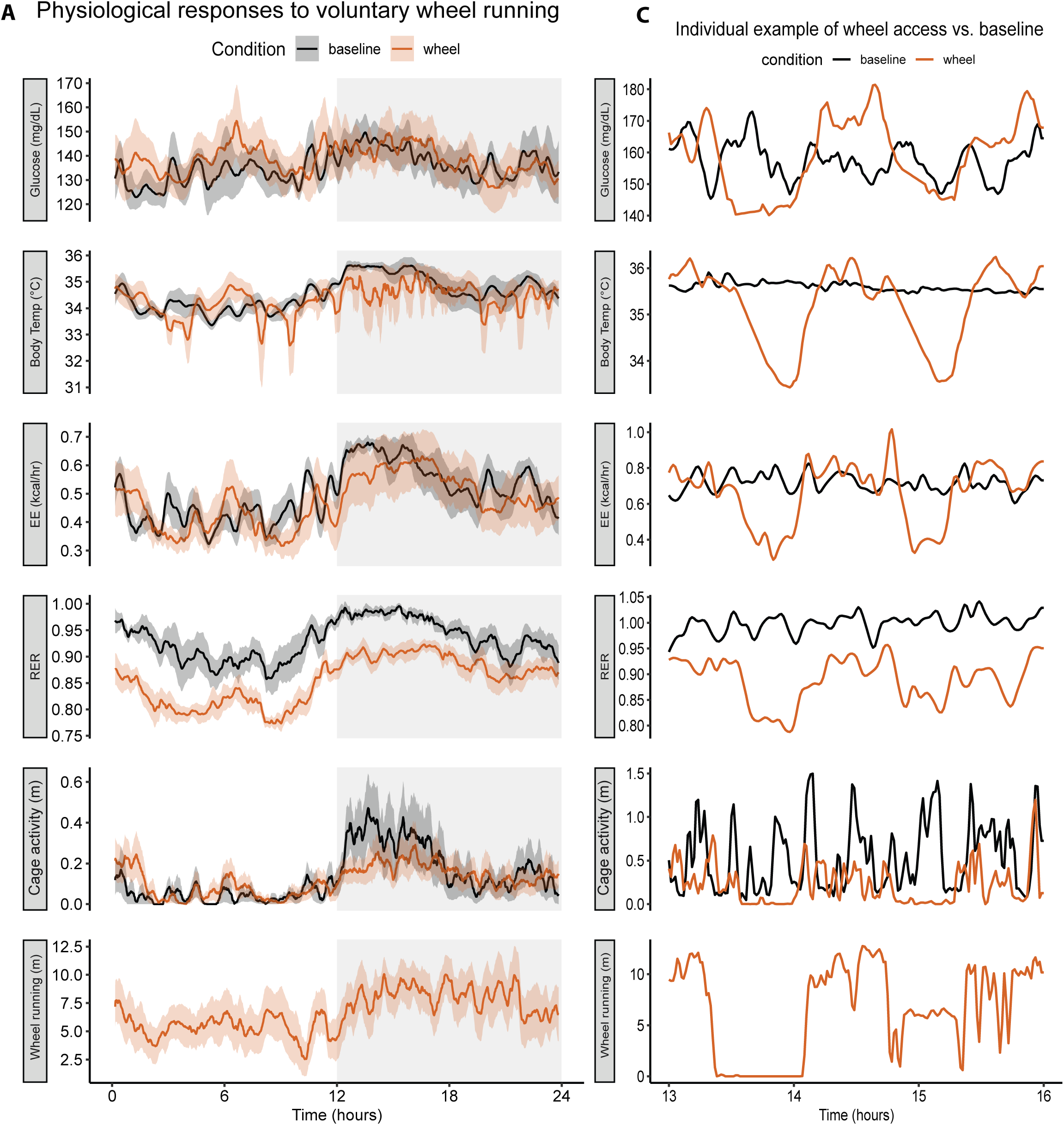
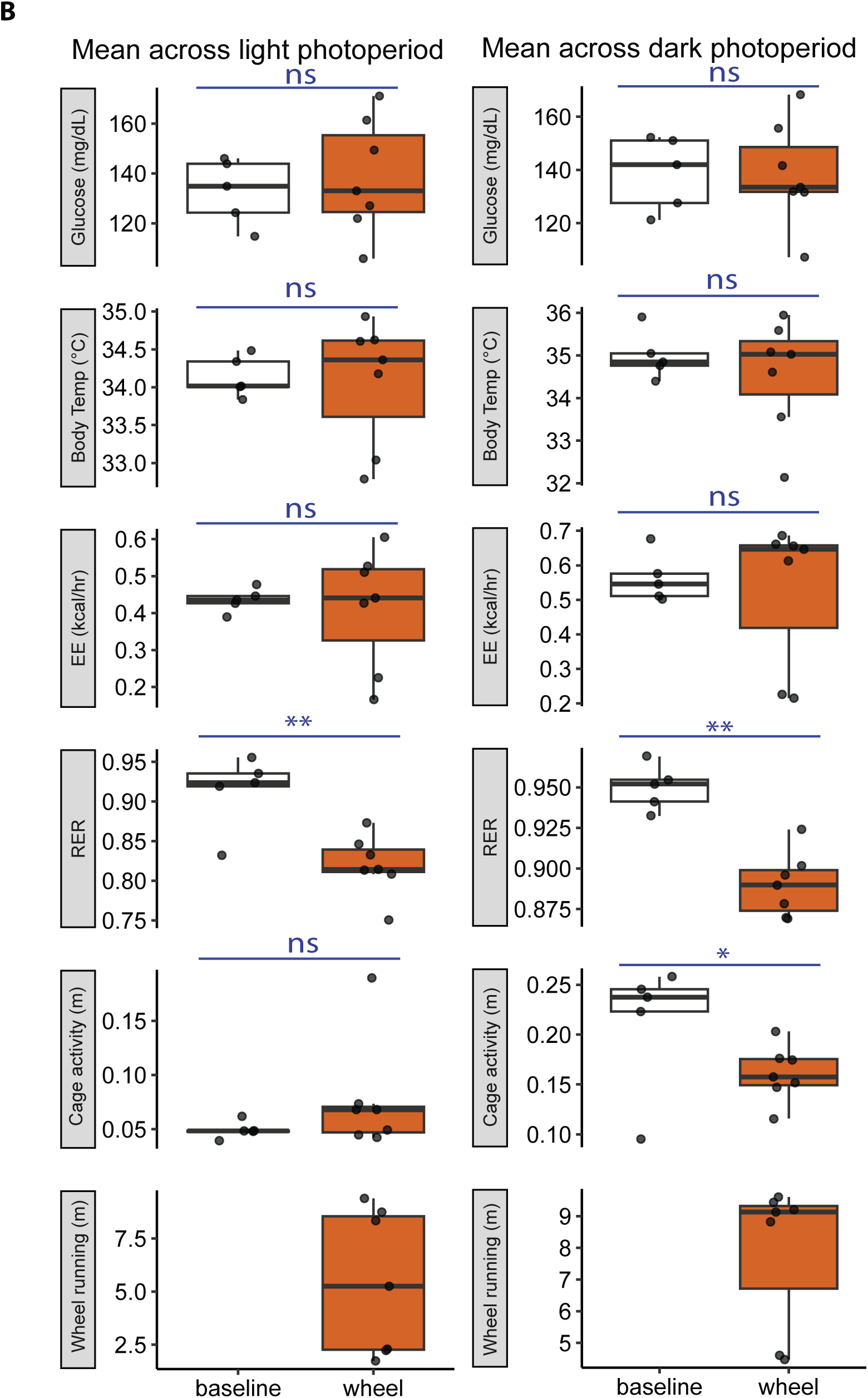
Metabolic effects of voluntary wheel running in chow-fed mice (baseline, n=5; wheel, n=7). **A)** Continuous recording of mice under baseline conditions and during access to a running wheel over 24h. Wheel-running group were introduced to wheels 1 day before the exper iment for brief acclimation. Gray shading indicates dark photoperiods. **B)** Boxplots showing mean values calculated across dark and light photoperiods. **C)** Recording of an individual mouse in dark photoperiod. Each point represents one mouse. Statistics by two-tailed unpaired t tests. ns, not significant; *p<0.05, **p<0.01, ***p<0.001, ****p<0.0001.

## Discussion

Responses to stress and metabolic challenges in mice are dynamic and context-dependent. In this study, we demonstrate that common procedures can themselves act as stressors that significantly alter metabolic readouts. For example, the transient rise in glucose at the initiation of fasting is counterintuitive and would likely be missed by conventional, low-frequency measurements. However, more frequent sampling is limited by the amount of blood loss that can be tolerated and, moreover, introduces additional stress that can, itself, alter the glucose levels being measured (8,17). To overcome these limitations, we employed continuous glucose monitoring (CGM) to provide high-resolution glucose measurements in mice and combined it with indirect calorimetry to enable real-time monitoring of multiple metabolic parameters simultaneously. Using this approach, we systematically examined the effects of common experimental variables, including fasting duration, handling, ambient temperature, and voluntary exercise, on glucose homeostasis and whole-body physiology.

Our results also underscore the impact of intervention by the investigator as a significant source of experimental variability. Procedures such as fasting and blood sampling introduce both environmental disturbance and physiological stress. For instance, removal of food containers can generate noise and disrupt the cage environment. This disturbance, combined with the abrupt absence of food, triggers an acute stress response, reflected by transient increases in glucose, body temperature, and locomotor activity. This stress-induced hyperthermia (18) is well appreciated in the neuroscience literature. In the context of GTT experiments, repeated tail blood sampling altered glucose dynamics, producing a more gradual decline following the peak compared to control mice. This suggests that handling-induced stress can impair glucose clearance, consistent with the known effects of stress hormones on glucose metabolism (4,8), and highlights a key advantage of CGM-based approaches that avoid repeated disturbance.

Environmental factors further modulated metabolic responses. Acute changes in ambient temperature produced rapid and coordinated shifts in glucose and energy balance, emphasizing the importance of controlling environmental conditions during metabolic phenotyping. As mice are typically housed at temperatures below thermoneutrality, they experience increased thermogenic demand, which influences energy expenditure and shifts substrate utilization (10,19). The cold-induced hyperglycemia in HFD mice is consistent with a mild-stress response and greater activation of the sympathetic nervous system and counterregulatory responses. Increased fatty acid oxidation can reduce glucose utilization (20), a factor which could contribute to hyperglycemia. However, most studies suggest higher total glucose oxidation during cold exposure along with increased hepatic glucose output (21). Thus, we hypothesize that hyperglycemia primarily reflects glucose mobilization in excess of uptake capacity, potentially exacerbated by insulin resistance in the HFD group. Testing this hypothesis will require direct measurements of glucose flux.

In addition, voluntary exercise induced transient increases in glucose, energy expenditure, body temperature, and respiratory exchange ratio (RER), reflecting acute metabolic activation. However, these changes did not translate into a sustained increase in total energy expenditure. Instead, RER was consistently reduced, indicating a shift toward increased fatty acid oxidation. Previous studies have shown that physical activity-induced energy expenditure (PAEE) can substitute for, rather than add to, thermogenic energy expenditure at room temperature (22,23). In this context, heat generated during exercise may be offset by compensatory reductions in other thermogenic processes, such as non-shivering thermogenesis, thereby masking increases in total energy expenditure during voluntary activity.

Combining CGM with indirect calorimetry brings together two different modalities for assessing physical activity and allows for direct comparison. The indirect calorimetry system uses a matrix of infrared sensors 1 cm apart and external to the cage to capture beam-breaks on the X-Y-Z axes. The number of beam breaks is used to infer physical activity. The CGM probes transmit radio signals to receivers placed beneath the cages, and changes to the intensity of the radio signal are used to calculate locomotor activity. We had assumed these measures would have a high degree of correlation. Indeed, the CGM radiofrequency-derived activity and calorimetry beam break activity were positively correlated, indicating that both systems capture general patterns of mouse movement. However, notable discrepancies were observed. A subset of time points exhibited high indirect calorimetry activity despite near-zero CGM activity (Figure S4, red points), whereas another subset showed elevated CGM activity in the absence of indirect calorimetry activity (Figure S4, green points). These discordant measurements reflect fundamental differences in how activity is quantified by the two systems. Consequently, low-amplitude behaviors such as grooming, rearing, or other cage interactions may contribute to indirect calorimetry activity but not CGM activity. Together, these findings indicate that CGM activity and indirect calorimetry activity measures capture related but distinct aspects of mouse behavior and should not be considered directly interchangeable. Given that indirect calorimetry activity integrates activity detected across the infrared beam array and captures a broader spectrum of spontaneous movement within the metabolic cage, we recommend the use of beam break-derived activity as the primary measure of locomotor activity in indirect calorimetry studies.

Two key technical considerations revealed by our study are the importance of maintaining accurate temporal precision in data alignment when dealing with high-resolution measurements and the importance of appropriate CGM calibration across the full physiological glucose range. With high-resolution measurements, recording the exact injection time of each individual animal for a GTT/ITT, or any test requiring staggered animal start times, and aligning both CGM and indirect calorimetry data together based on start time is essential to capture these dynamic metabolic responses accurately at a high resolution. We also found that accurate measurements, particularly at hypoglycemic glucose levels and in long-duration studies, required more frequent multipoint calibrations and additional multipoint calibrations covering the lower dynamic range of blood glucose, both of which are beyond standard protocols provided by the manufacturer (24). Specifically, incorporating low glucose level calibration points derived from the ITT improved accuracy in the hypoglycemic range, which is not well captured by the recommended hyperglycemic GTT-based calibrations. We also observed improved agreement between CGM and glucometer readings when both additional calibration points were included and more frequent multipoint calibrations were performed, likely due to correction of sensor drift over time, a known limitation of CGM systems (25). Together, these results suggest that accurate temporal resolution should be maintained to align high-resolution measurements, CGM probes should be calibrated within the expected glucose range of the experiment and that repeated calibrations, both single-point and multi-point, are important for maintaining accuracy, especially in longer-duration studies.

Collectively, our findings demonstrate that experimental conditions, including fasting duration, handling, environmental temperature, and physical activity, can profoundly influence metabolic measurements in mice. These results underscore the importance of minimizing handling and standardizing environmental variables when assessing glucose metabolism in vivo. More broadly, our study highlights the value of combining high-resolution approaches such as CGM and indirect calorimetry to capture the true dynamics of metabolic regulation.

### Limitations

Although CGM enables high resolution and minimizes handling-related artifacts, its accuracy depends on the accuracy of the surgery and appropriate calibration. CGM implantation requires major surgery involving insertion of the probe through the carotid artery with advancement into the aortic arch, a technically challenging procedure that requires a skilled surgeon to execute accurately and minimize surgical mortality. This invasive procedure may also introduce baseline physiological changes despite adequate recovery periods. In addition, despite incorporating additional multipoint calibrations, sensor drift may still affect absolute glucose values over time. Furthermore, the probes are single-use and relatively expensive (approximately $1,100 per probe), potentially restricting scalability and broader adoption in large cohort studies. Our study was also performed under controlled laboratory conditions using a defined mouse cohort, and metabolic responses may differ across strains, sexes, ages, or physiological states, potentially limiting generalizability.

## Methods

### Animals and housing

All animal experiments were approved by the Institutional Animal Care and Use Committee (IACUC) at Beth Israel Deaconess Medical Center and the University of Pennsylvania. Male C57BL/6J mice (14-20 weeks old) were obtained from The Jackson Laboratory and maintained on either a standard chow diet or a 60% high-fat diet, as indicated. Mice were maintained under a 12 h light/12 h dark cycle (06:00-18:00 or 07:00-19:00) at ∼22-23 °C with ad libitum access to food and water unless otherwise specified. Animals were acclimated to experimental conditions prior to data collection.

### Continuous glucose monitoring

Continuous glucose monitoring (CGM) was performed using HD-XG telemetry probes (Data Sciences International), which measure blood glucose directly from the aortic arch and core or subcutaneous body temperature from either the peritoneal cavity or subcutaneous site, respectively. Probes were surgically implanted according to manufacturer guidelines (26), and mice were allowed to recover for 7-10 days before data collection.

Probes were calibrated using a combination of multipoint and single-point calibrations based on glucometer measurements from tail blood samples. To ensure accurate measurements across a wide dynamic range, additional multipoint calibrations were performed beyond standard GTT conditions, including calibrations in lower glucose ranges using insulin tolerance test (ITT) conditions. Calibration points were repeated if measurements differed by more than 10%.

### Indirect calorimetry

Metabolic measurements were performed using the Promethion indirect calorimetry system (Sable Systems International). Mice were individually housed in metabolic cages with continuous access to food (Labdiet 5008 or 5010, metabolizable energy 3.19 or 3.00 kcal/g, respectively, or Research Diets D12492i, 60% kcal from fat, 5.21 kcal/g) and water unless otherwise specified. Oxygen consumption (VO₂) and carbon dioxide production (VCO₂) were measured every 2-3 minutes and used to calculate energy expenditure (EE) using the Weir equation. Respiratory exchange ratio (RER) was calculated as VCO₂/VO₂. Cage activity was measured as the 1-minute change in the cumulative AllMeters parameter generated from the Promethion X-Y-Z infrared beam-break array. Telemetry receiver plates were placed beneath the cages to enable simultaneous collection of CGM and physiological data, and metal dividers were placed between cages to prevent radiofrequency interference between probes. All data streams were aligned and imputed to a 1-minute resolution based on system timestamps to match glucose and body temperature recordings by CGM probes.

### Experimental interventions

Fasting was initiated by removing food containers or closing food access control doors (Sable Systems International) with a cage change to woodchip or ALPHA-dri bedding (Shepherd Specialty Papers) to minimize residual food. Mice retained access to water throughout fasting. Refeeding was performed by returning food to the cage or opening food access control doors. Cage changes and food removal were associated with brief data dropout, which was accounted for during data interpretation.

Glucose tolerance tests (GTT) were performed following a 5-hour fast, with glucose administered as a sterile-filtered 20% solution intraperitoneally at a dose of 2 g/kg body weight. Blood glucose measurements were obtained via tail-tip sampling using a handheld glucometer. For CGM-based analyses, continuous glucose data were used to assess glucose dynamics without repeated handling. Insulin tolerance tests (ITT) were performed with insulin administered intraperitoneally at 0.75 U/kg body weight, following fasting periods that varied depending on the experimental design.

Ambient temperature experiments were conducted using a temperature-controlled cabinet integrated with the Promethion system, allowing assessment of metabolic responses under controlled thermal conditions.

Voluntary exercise was assessed by providing mice with free access to a running wheel within the metabolic cages. Wheel-running activity was recorded alongside other physiological parameters.

### Data processing and analysis

Data were analyzed in R 4.4.3 using the tidyverse package. CGM and indirect calorimetry data were synchronized based on timestamps and smoothed for visualization using rolling mean filters (typically 5-20 minutes). CGM data alignment was performed using a custom R/Shiny pipeline. The application imported all CGM and indirect calorimetry files from a selected directory and processed them in sorted order. A metadata table was required to define file-to-sample mapping (file_index), cage assignment (cage_num), and sample annotations (sample_name, condition). Metadata start times were standardized and parsed using multiformat rules. For each sample, the algorithm identified the row corresponding to the metadata start time (minute-level match with nearest-time fallback). This row was defined as time 0, corresponding to the individual injection times for each mouse. The program then extracted a fixed baseline window before time 0 and a fixed experimental window after time 0, based on user-defined durations. Cage-specific variables were mapped to standardized metric names and merged into a single long-format dataset with relative time (Minutes) and sample/group labels for downstream analysis. Missing values due to telemetry signal dropout were interpolated using the zoo package (27).

Glucose responses during GTTs and ITTs were quantified by calculating the area of the curve (AOC) for the glucose excursion relative to each animal’s baseline glucose concentration using trapezoidal integration. This approach minimizes the influence of differences in baseline glycemia between animals and experimental groups and provides a measure of the magnitude of the glucose response to the metabolic challenge. The same analytical framework was applied to both GTTs and ITTs to enable consistent comparison of glycemic excursions across different physiological interventions, as recommended previously (28). Group comparisons were performed using ANOVA followed by appropriate post hoc tests. Statistical significance was defined as p < 0.05.

## Acknowledgements

This work is supported by NIH grants RC2DK142612, P30DK135043, the Boston Area Diabetes and Endocrinology Research Center to ASB. Rodent Metabolic Phenotyping Core (RRID: SCR_022427) is supported in part by National Institutes of Health (NIH) grant S10-OD025098 to JAB, the Cox Institute, and the Institute for Diabetes, Obesity and Metabolism (Perelman School of Medicine, University of Pennsylvania). This work is dedicated to the memory of Dr. Jeffrey Ishibashi for his unwavering support, innovative techniques, and most importantly, his tireless effort to understand and advance all aspects of science.

---

**Supplementary Figure 1.**
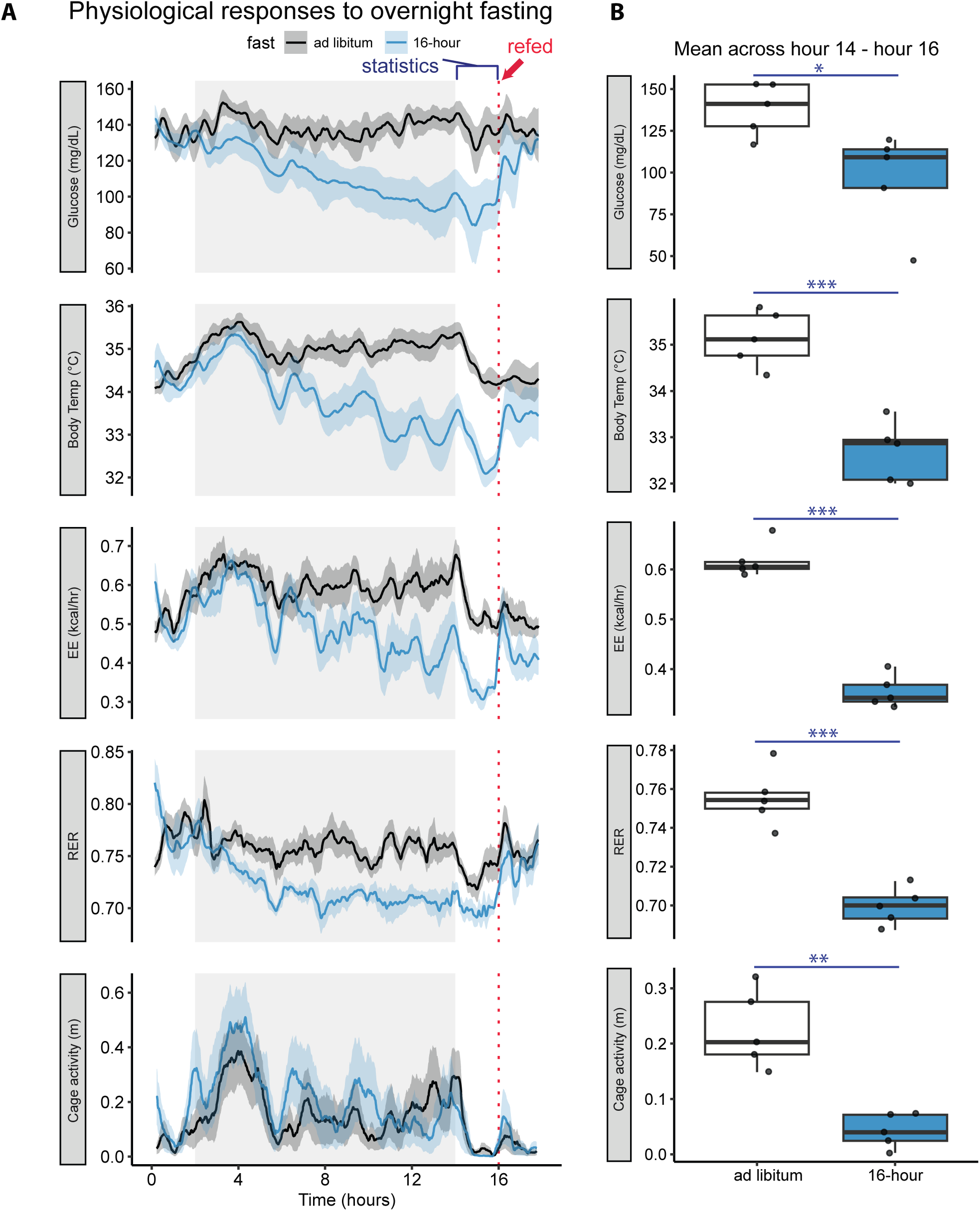
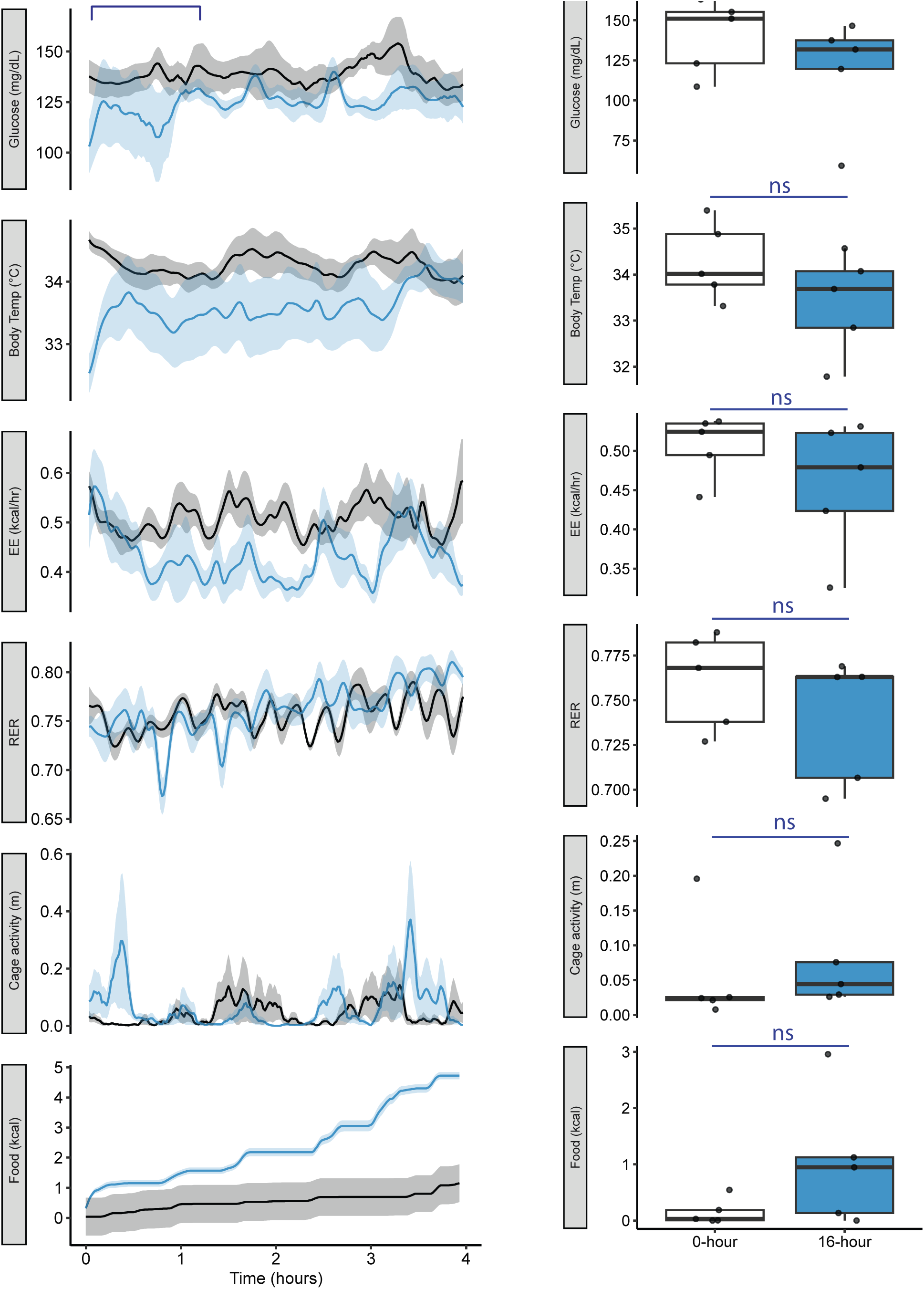
Effects of overnight fasting and refeeding on metabolic parameters in HFD-fed mice (n=5). **A)** Recordings of HFD-fed mice during 16h overnight fasting. Solid lines represent the mean value at each minute, shaded areas represent SEM. Grey shadings indicate dark photoperiods. **B)** Boxplots showing mean values calculated over hours 14–16. Each point represents one mouse. Statistics by two-tailed unpaired t tests. **C)** Recordings of refeeding following overnight fast, mice were refed at t = 0. Solid lines represent the mean value at each minute, shaded areas represent SEM. **D)** Boxplots showing mean values calculated over hours 0-1 after refeeding. Each point represents one mouse. Statistics by two-tailed unpaired t tests. *p<0.05, **p<0.01, ***p<0.001, ****p<0.0001.

**Supplementary Figure 2.**
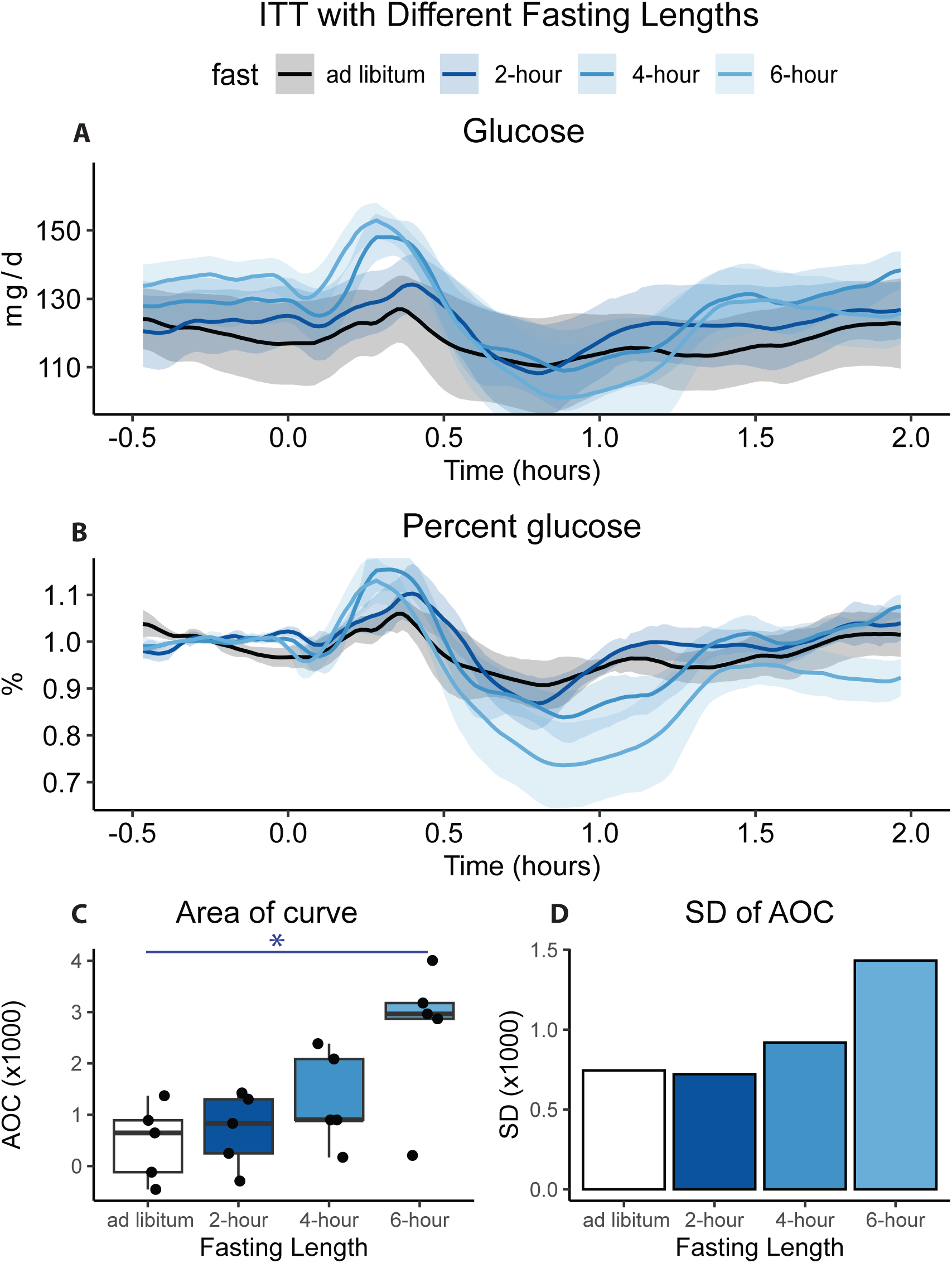
Effects of fasting duration on insulin tolerance test in HFD-fed mice (n = 5). **A)** Continuous glucose monitoring during ITTs plotted in absolute experimental time. Mice underwent 0-, 2-, 4-, or 6-h fasting at the time of insulin injection (t = 0). **B)** Glucose values expressed as percent of baseline for the same recordings. Baseline glucose was defined as the average glucose during the 1 h preceding insulin injection. **C)** Area over the curve (AOC) calculated from percent glucose traces. Each point represents one mouse. Statistical significance was determined by one-way ANOVA; *p<0.05. **D)** Standard deviation (SD) of AOC across mice for each fasting duration.

**Supplementary Figure 3.**
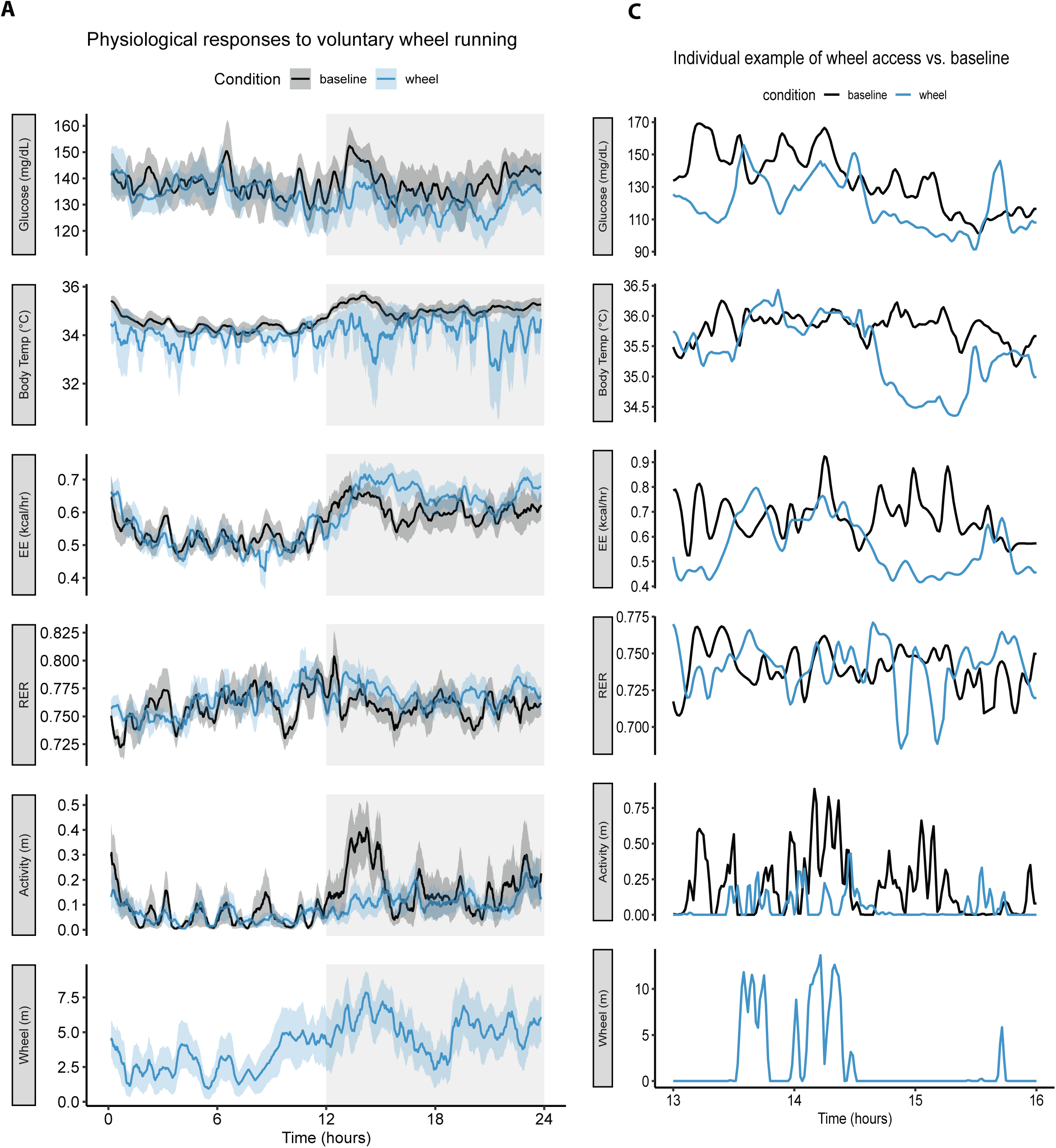
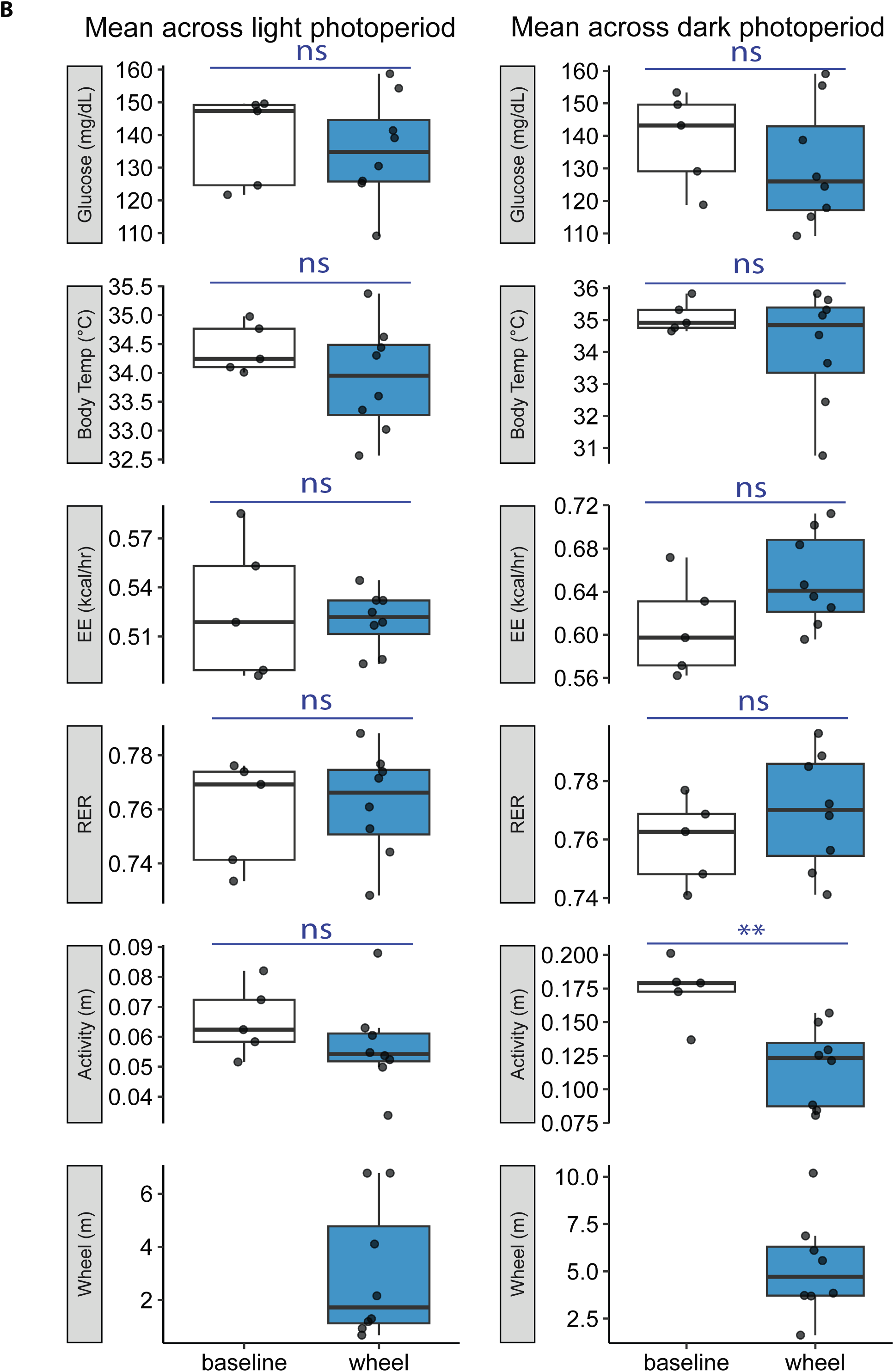
Metabolic effects of voluntary wheel running in HFD-fed mice (baseline, n=5; wheel, n=7). **A)** Continuous recording of mice under baseline conditions and during access to a running wheel over 24h. Wheel-running group were introduced to wheels 1 day before the experiment for brief acclimation. Gray shading indicates dark photoperiods. **B)** Boxplots showing mean values calculated across dark and light photoperiods. **C)** Recording of an individual mouse in dark photoperiod. Each point represents one mouse. Statistics by two-tailed unpaired t tests. ns, not significant; **p<0.01.

**Supplementary Figure 4.**
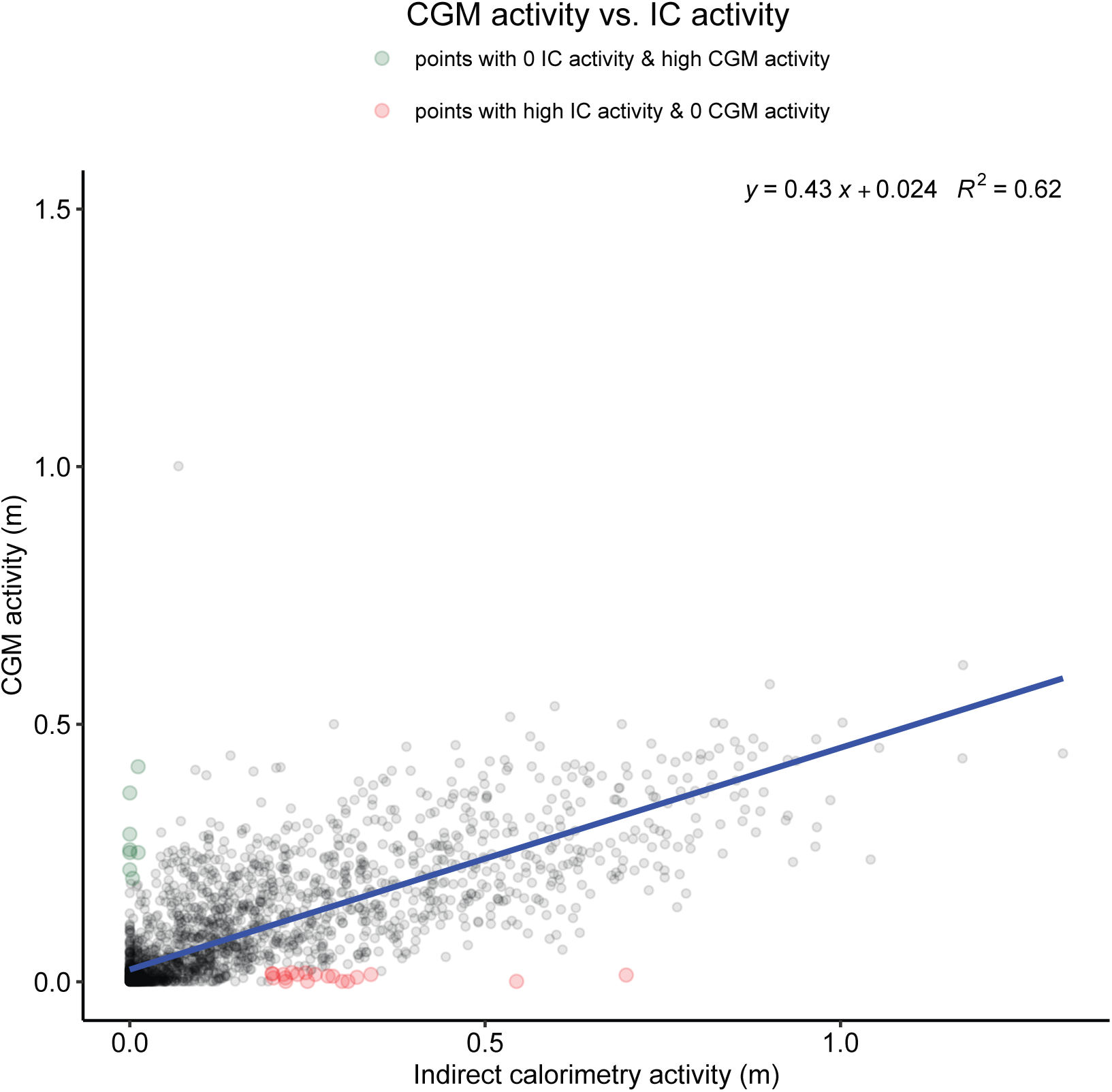
CGM radiofrequency activity vs. Indirect calorimetry beam-break activity. All time points collected during a 24-hour baseline recordings, y-axis shows the CGM-derived activity, x-axis shows the IC activity derived from Allmeters. Red points denote observations with high IC activity but minimal CGM activity, whereas green points denote observations with elevated CGM activity despite no IC activity. The blue line represents the linear regression fit.

